# A Computer Vision Dataset for Pollinator Detection under Real Field Conditions

**DOI:** 10.1101/2025.10.27.682286

**Authors:** Yue Linn Chong, Phillip Nachtweide, Julian Bauer, Andrée Hamm, Jana Kierdorf, Lukas Drees, Cyrill Stachniss, Jens Behley, Thomas F. Döring, Ribana Roscher, Sabine J. Seidel, Antonia Veronika Mayr

## Abstract

The widespread decline in biodiversity and abundance of pollinator insects is expected to provoke cascading effects on food security and jeopardize ecosystem services crucial for many crops and wild plants. Pollinator monitoring is a crucial element in preventing further decline of pollinators, to which computer vision approaches can make essential contributions. To facilitate research in such approaches, we present a dataset for pollinator detection with accurate annotations. We develop the dataset with an iterative semi-automatic annotation approach, which leverages YOLO to assist with human annotation. We quantify the impact of multiple levels of errors in annotations on training and report the increase in mAP of 28.7% at the final iteration when compared to the manual annotations. Our dataset encompasses pollinator detection for honeybees and bumblebees across various flower treatments over multiple days. Our dataset facilitates the development of deep learning-based methods for automatic large-scale pollinator detection under various real-world field conditions, as well as adjacent computer vision tasks such as small object detection and label correction.

## 1 Introduction

There is a widespread decline in biodiversity and abundance of pollinators, partly due to agricultural intensification [5,6,7,33]. This decline may provoke cascading effects on ecosystem services [30], such as pollination, which is crucial for many crops and wild flowering plants [4,13]. In preventing further decline of pollinators, pollinator monitoring is crucial, providing measurements on their occurrence to assess conservation strategies such as flower enrichment intercropping in agricultural fields. Traditional insect monitoring methods include trapping, active visual surveys, and other highly manual methods [18], which require intensive manual labor and domain expertise, thus they are unsuitable for large-scale monitoring. Previous works [3, 14, 22, 23, 28] propose replacing the traditional methods with automatic deep learning-based methods. To this end, they manually annotate a dataset for training the deep learning models. However, manual annotations often contain human errors. In particular, annotating pollinators while they are foraging in real fields can be challenging due to the small size of the pollinators and the cluttered background, see Fig. 1 and Fig. 3 for examples. This challenge results in incomplete annotations, that is, where the annotations miss some pollinators, yielding annotations with a lower recall. These erroneous annotations can lead to poor model performance.

**Fig. 1.**
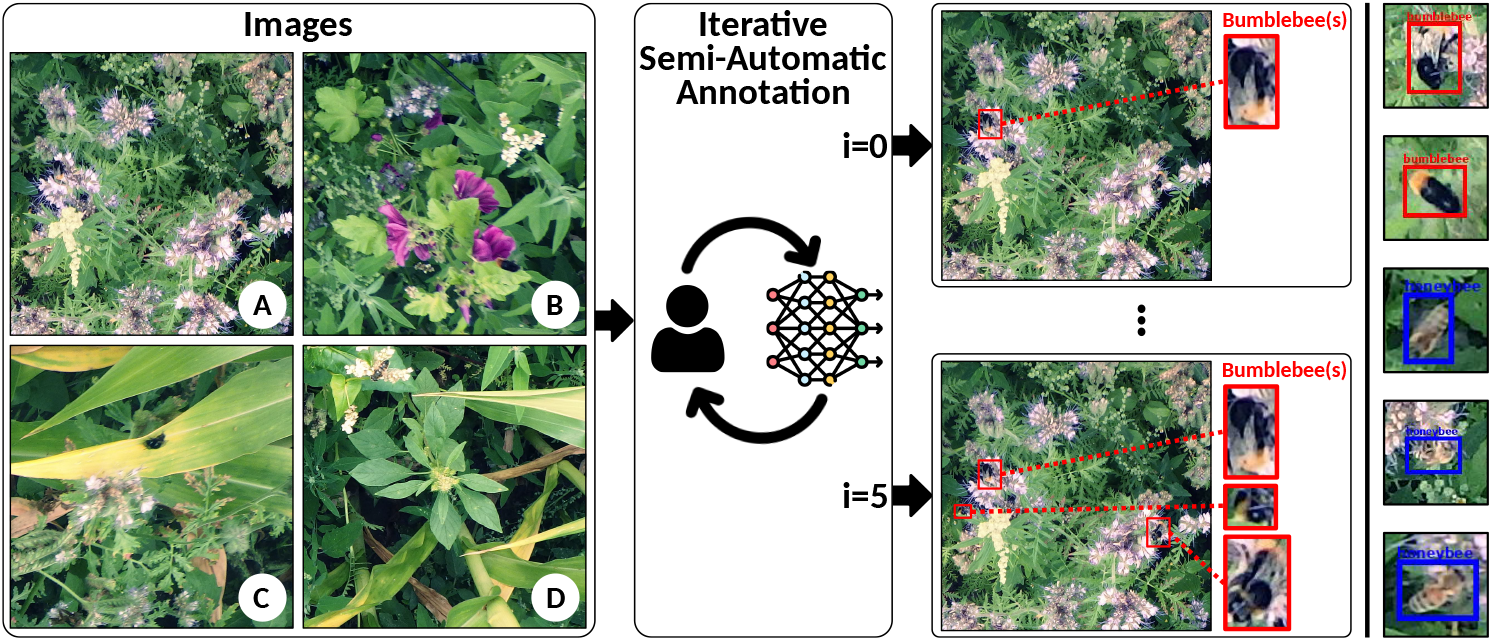
Our dataset consists of images we collect at various treatments of (A) phacelia, (B) wildflowers, (C) phacelia-maize, and (D) wildflower-maize treatments. We annotate the images using an iterative semi-automatic approach that leverages deep learning. With each iteration, we obtain a set of annotations with decreasing levels of errors, finally yielding accurate annotations. On the right, we show examples of some bumble-bees (red) and honeybees (blue).

To tackle this challenge, we record new data and annotate it using an iterative, semi-automatic approach to correct annotations, aiming to annotate all pollinators in a set of images. Our approach iteratively improves the annotations by leveraging a deep learning model, which we train in tandem with annotating the dataset. We apply our annotation approach to high-resolution RGB images we collect from a field trial with flower treatments of phacelia, mixed wildflowers, and flower-enriched maize intercrops over several days, and annotate honeybees (*Apis mellifera*), bumblebees (*Bombus* spp.), and other pollinators. Fig. 1 illustrates our dataset development process. With the dataset, we train and evaluate several baseline object detection methods.

We provide multiple sets of annotations with varying error levels which we generate from the intermediate iterations of the annotation approach, where in which the errors decrease with each iteration. These sets of annotations with human errors enable studies of model performance at incremental degrees, towards increasing method robustness, that is especially relevant in highly specialized domains such as entomology [32] and agriculture [26, 31] where annotators’ expertise is critical. To facilitate research in this direction, we evaluate the model performance when trained with these intermediate iterations’ annotations.

In summary, our three main contributions are: (i) We present an open-source dataset for pollinator detection from images we recorded from our field trial with complete and accurate annotations, on which evaluate object detection baselines; (ii) We present an iterative semi-automatic approach for annotation; (iii) We present several sets of annotations with varying levels of errors and quantify the drop in model performance with each level.

## 2 Related Work

Deep learning methods for pollinator monitoring have gotten increasing attention [1, 3, 10, 14, 22, 23] with several works also publishing annotated datasets, as summarized in Tab. 1. While most datasets focus on honeybees [14, 19, 23, 27], our dataset includes annotations for both honeybees and bumblebees, as well as annotations for species with fewer instances under the class “other insect”. In contrast to the datasets of beehives [19, 27] with structured static backgrounds, our dataset provides annotated data under real field conditions for real-world pollinator monitoring where the background of plants is often cluttered.

**Table 1.**
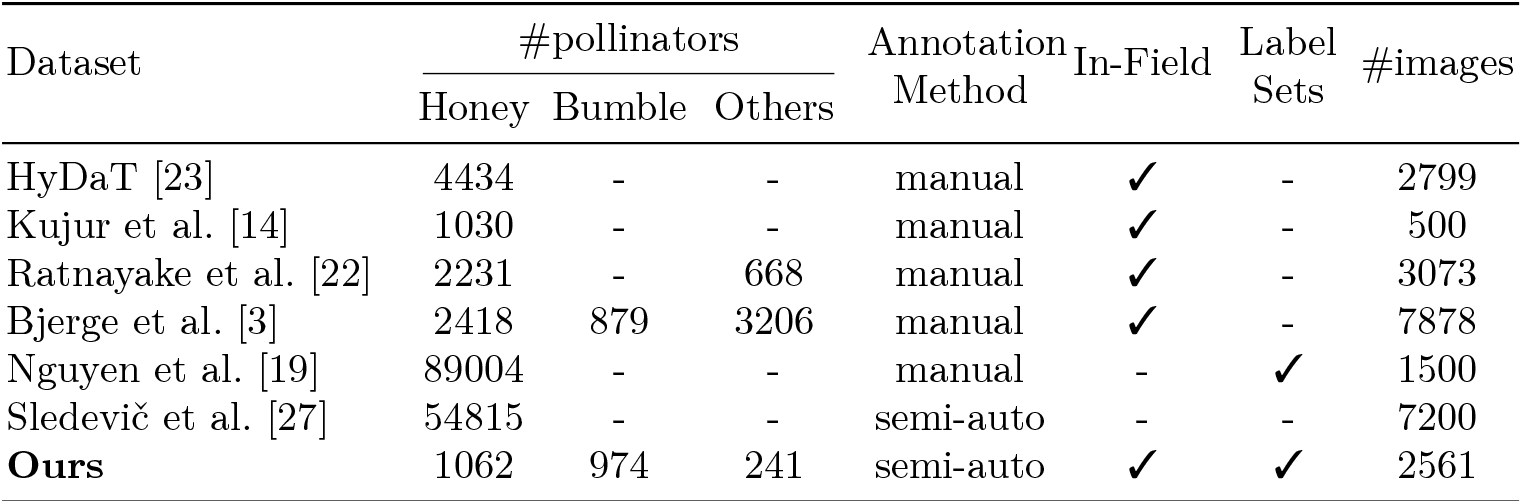
Comparison of datasets in the pollinator monitoring domain. “*✓*” denotes that the characteristics are present in the dataset, and “-” denotes that the characteristic is not present in the dataset. Label sets indicate if the dataset also provides noisy annotations in addition to perfect annotations.

| Dataset | #pollinators |  |  | Annotation Method | In-Field | Label Sets | #images |
| --- | --- | --- | --- | --- | --- | --- | --- |
|  | Honey | Bumble | Others |  |  |  |  |
| HyDaT [23] | 4434 | - | - | manual | ✓ | - | 2799 |
| Kujur et al. [14] | 1030 | - | - | manual | ✓ | - | 500 |
| Ratnayake et al. [22] | 2231 | - | 668 | manual | ✓ | - | 3073 |
| Bjerge et al. [3] | 2418 | 879 | 3206 | manual | ✓ | - | 7878 |
| Nguyen et al. [19] | 89004 | - | - | manual | - | ✓ | 1500 |
| Sledevič et al. [27] | 54815 | - | - | semi-auto | - | - | 7200 |
| **Ours** | 1062 | 974 | 241 | semi-auto | ✓ | ✓ | 2561 |

Most datasets are manually annotated [3, 14, 19, 22, 23]. However, manual annotations are prone to human errors, so several works propose assisting manual annotation with machine learning in adjacent domains, such as slide cell counting [2, 17] and coral reef segmentation [21]. We follow a similar approach, concurrently developing the annotations and a deep learning model. Specifically, we adopt a similar approach to Bertram et al. [2], where the deep learning model proposes new annotations for human annotators to verify. Unlike Bertram et al., we iterate the model training and manual annotations, generating a set of annotations at each iteration to obtain accurate annotations.

In pollinator monitoring, Sledevi*č* and Matuzevi*č*ius [27] propose using deep learning for semi-automatic labeling. However, their approach directly uses manually corrected predictions as the final annotations. Conversely, our approach iteratively builds the dataset annotation while training the model to achieve more accurate annotations under real-world field conditions. Although our dataset is not the largest, we offer a diverse range of images spanning several days and various plant treatments.

Erroneous annotations can lead to poor method performance, but errors in annotations are often inevitable. Several methods claim to be more robust against noisy training annotations [15, 20, 29]. Concurrently, new datasets [8, 9] are also available for assessing the method robustness or label correction. In pollinator monitoring, Nguyen et al. [19] proposes a method for handling noisy labels for bee counting. However, they synthetically generate a noisy dataset by injecting noise into accurate annotations. In contrast, our dataset provides several sets of annotations with varying completeness levels, with noise resulting from human error, which may not be replicable by adding random noise.

In summary, we present a dataset with accurate annotations for detecting honeybees, bumblebees, and other insects under real-world field conditions. Using an iterative semi-automatic annotation approach, we generate accurate annotations along with multiple intermediate sets of annotations, each with different levels of errors. The dataset can support future work on model robustness, label correction, and the detection of small objects.

## 3 Dataset and Approach

### 3.1 Our Iterative Semi-Automatic Data Annotation Approach

The iterative semi-automatic annotation approach aims to improve the accuracy of the annotations with each iteration, mainly to ensure that no pollinators are left out (i.e., not annotated). In particular, we focus on overcoming the challenges of annotating all small pollinators and ensuring that our annotations are complete, as pollinator counts are used for downstream pollinator monitoring analysis. Fig. 2 illustrates the overview of the approach.

**Fig. 2.**
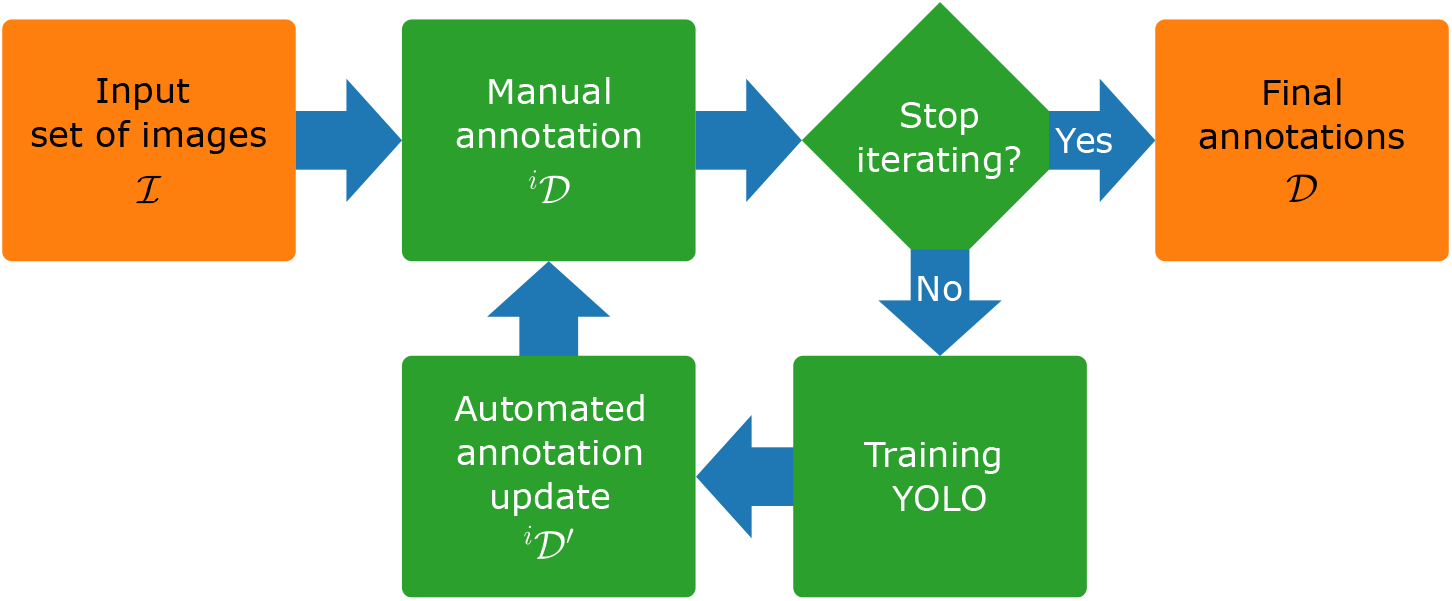
Overview of the iterative semi-automatic annotation approach. The approach takes as input a set of images ℐ. From ℐ, we use manual annotations as the initial annotations and train YOLO with them. We then use YOLO predictions on ℐ to automatically update the annotations. Human annotators then review the annotations, and we repeat the process until the count of pollinators no longer increases. We use the annotations from the last iteration as the final annotations for our dataset.

The approach takes in as input a set of RGB images, ℐ= *{***I**_1_, **I**_2_, …, **I**_*N*_*}* and for each **I**_*n*_, outputs corresponding detections

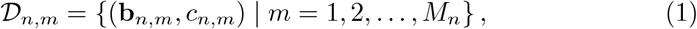

where **b**_*n,m*_ = [*x*_*n,m*_, *y*_*n,m*_, *w*_*n,m*_, *h*_*n,m*_] is the parameterized bounding box centered at (*x*_*n,m*_, *y*_*n,m*_) with width *w*_*n,m*_ and height *h*_*n,m*_, and corresponding object class *c*_*n,m*_ ∈ {honeybee, bumblebee, others *}*. We use the images we collect as described in Section 3.2 as inputs. The approach is semi-automatic, as we train a YOLO [11] model in each iteration and run inference with the YOLO to obtain predictions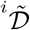. We then manually review and correct these annotations; and repeat these steps until we satisfy the stopping criterion. The following subsections elaborate on the modules within our approach.

#### Manual Annotation

As initialization (at iteration *i* = 0), for each image **I**_*n*_, the human annotators manually annotate the bounding boxes ^0^**b**_*n,m*_ and classes ^0^*c*_*n,m*_ of each pollinator individual. Note that at the earlier iterations, we expect these manual annotations to contain errors, including false negatives, i.e., with missed annotations. In subsequent iterations, in addition to the images, the human annotators are given prior annotations for image **I**_*n*_

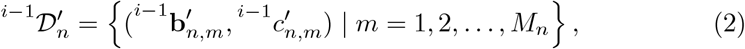

which they then correct, paying careful attention to when the annotations disagree with the previous iteration. The annotators verify the location and class of each new prediction, improving both precision and recall. The annotators locate all target pollinators within a reasonable timeframe and can remove or add annotations freely. Note that locating all pollinators in images is challenging, as the pollinators are small, and the background is crowded with plants. We denote the annotations the human annotators output as ^*i*^ 𝒟_*n*_, i.e., the annotations for the current iteration *i*.

#### Stopping Criterion

Toward obtaining the complete annotations, we repeat our process until the number of pollinator annotations no longer increases from the previous iteration |^*i*^ 𝒟_*n*_| ≤ | ^*i*−1^𝒟 _*n*_|. We reach this criterion after the fifth iteration, so we finalize the annotations at *i* = 5 and form the final dataset.

#### Training YOLO

To assist with manual annotation, we train a YOLO [11] model to automatically detect pollinators and help correct errors in the manual annotation. To this end, we train YOLO with the possibly erroneous annotations at the current iteration ^*i*^ 𝒟_*n*_. Several types of errors exist in annotations, including false negatives from missing annotations and misclassifications. For our pollinator monitoring use case, detecting missing annotations is crucial, as pollinators are small and easily overlooked by human annotators among the flowers in the background. To mitigate the effects of missed annotations which can incorrectly penalize YOLO during training, we crop each image to include only the areas that the annotators have manually checked, in hopes to reduce the number of missing pollinators. Since the human annotators review the annotations from the previous iteration 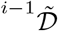, we patch the areas around these annotations and do not patch areas with no prior detections.

Specifically, for each image **I**_*n*_ and corresponding latest manually corrected annotations ^*i*^ 𝒟_*n*_, we obtain a set of smaller images Î = *{*Î_*n,m*_*}*, with Î_*n,m*_ cropped from *I*_*n*_ centered around (*x*_*m*_ + *δ*_*x*_, *y*_*m*_ + *δ*_*y*_), where *δ*_∗_ is a random perturbation to avoid centering the image around each annotation in 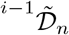. Each Î_*n,m*_ has the size of *λ*_size_ *× λ*_size_ pixels. We use *λ*_size_ = 1024 in our experiments and provide an ablation study below. Correspondingly, we translate ^*i*^𝒟_*n*_ into the frame of each Î_*n,m*_ to obtain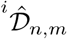, and we train YOLO with the set of inputs and labels (Î _*n,m*_,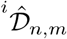). Fig. 3 illustrates this cropping process. Furthermore, to reduce misclassification, we introduce the class “other insects” for when the classification between honeybee and bumblebee is unclear to the human annotators. The annotators check where YOLO disagrees with the annotated class and reassign annotations to “other insects” if they cannot clearly distinguish between honeybee and bumblebee. The annotators also adjust the size of the bounding boxes to fit the pollinators as tight as possible while encapsulating the whole pollinator.

In the first iteration, we initialize the model weights with weights pre-trained with COCO [16]. From the second iteration onward, we initialize the YOLO weights with the weights from the first iteration. We did not use the weights from later iterations for initialization, as we found that this led to poorer performance in our preliminary experiments, possibly because the network is stuck in a local minimum from previous iterations. We use the medium-sized YOLO architecture and train it with a linear learning rate scheduler from 0.01 to 0.1 and stochastic gradient descent, a batch size of 6, early stopping, and a maximum of 600 epochs, with the image size 5120 pixels. Furthermore, we apply augmentations of color jittering in the HSV space, translation, scaling, flipping, mosaic, and mix-up.

#### Automated Annotation Update

After training, we ran inference with YOLO to detect target pollinators in all images in ℐ to obtain predictions

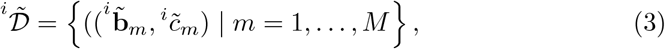

where we mark 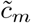 with a different color such that the human annotators can visually differentiate between new YOLO predictions and manual annotations current iteration via the annotation tool graphical user interface. We also mark the individuals that are the same between 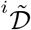and ^*i*^𝒟, where 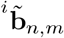 and ^*i*^**b**_*n,m*_ satisfy the intersection-over-union (IoU) ≥ 0.7 and 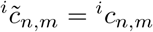. During inference, we set the hyperparameters to enable a higher detection rate of pollinators, specifically by setting the confidence threshold to a lower value of 0.25, the IoU threshold of 0.45, and the maximum number of detections of 1000. Finally, we combine the YOLO detections with those of the current manual annotations,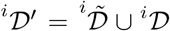. Note that this stage does not remove any annotations but instead aims to prepare the data for manual checks of all annotations, including “true positive” cases, as current annotations may also be incorrect.

#### Ablations

In the semi-automatic annotation approach, we cropped out smaller images Î_*n*_ from the entire image **I**_*n*_ to exclude areas where human annotators have not explicitly verified, thereby preventing training with erroneous data. In this section, we evaluate this design choice by comparing the number of annotations we incorrectly missed with our patching approach to using random patching or the whole image instead. We use the labels from the initial manual annotation (*i* = 0) as incorrect labels since this set of annotations has the highest number of missing bees. Tab. 2 shows the number of individuals missed in each training data preparation. If we train with the whole image, the ratio of the missed bees to the present bees would be high compared to our patching approach, which misses fewer bees. Random patching of the images is not suitable, as the training data contains only a few bees and has a large proportion of missing bees. Thus, our approach in patching for preparing the training data is the most suitable for developing our dataset.

**Table 2.** Ablation study on different preprocessing methods for training YOLO, showing the total number of annotations correctly present versus incorrectly missed in the training data when using manual annotation only (*i* = 0).

| Method | #correct | #missed |
| --- | --- | --- |
| Whole image | 646 | 847 |
| Random | 51 | 81 |
| **Ours** | 840 | 201 |

### 3.2 Data Collection

We conduct a field experiment with flower treatments for data collection, shown in Fig. 4. We set up RGB cameras to capture images of the field and monitor pollinators under real field conditions. The target pollinators are honeybees and bumblebees, where we assign the “honeybee” class to *Apis mellifera* individuals and “bumblebee” to individuals of the genus *Bombus* spp. The “other insects” class is the catch-all class for insects that human annotators cannot definitively identify by visual inspection as either a “honeybee” or a “bumblebee”, and also includes other bee mimic species such as *Eristalis tenax* or *Episyrphus balteatus*.

**Fig. 3.**
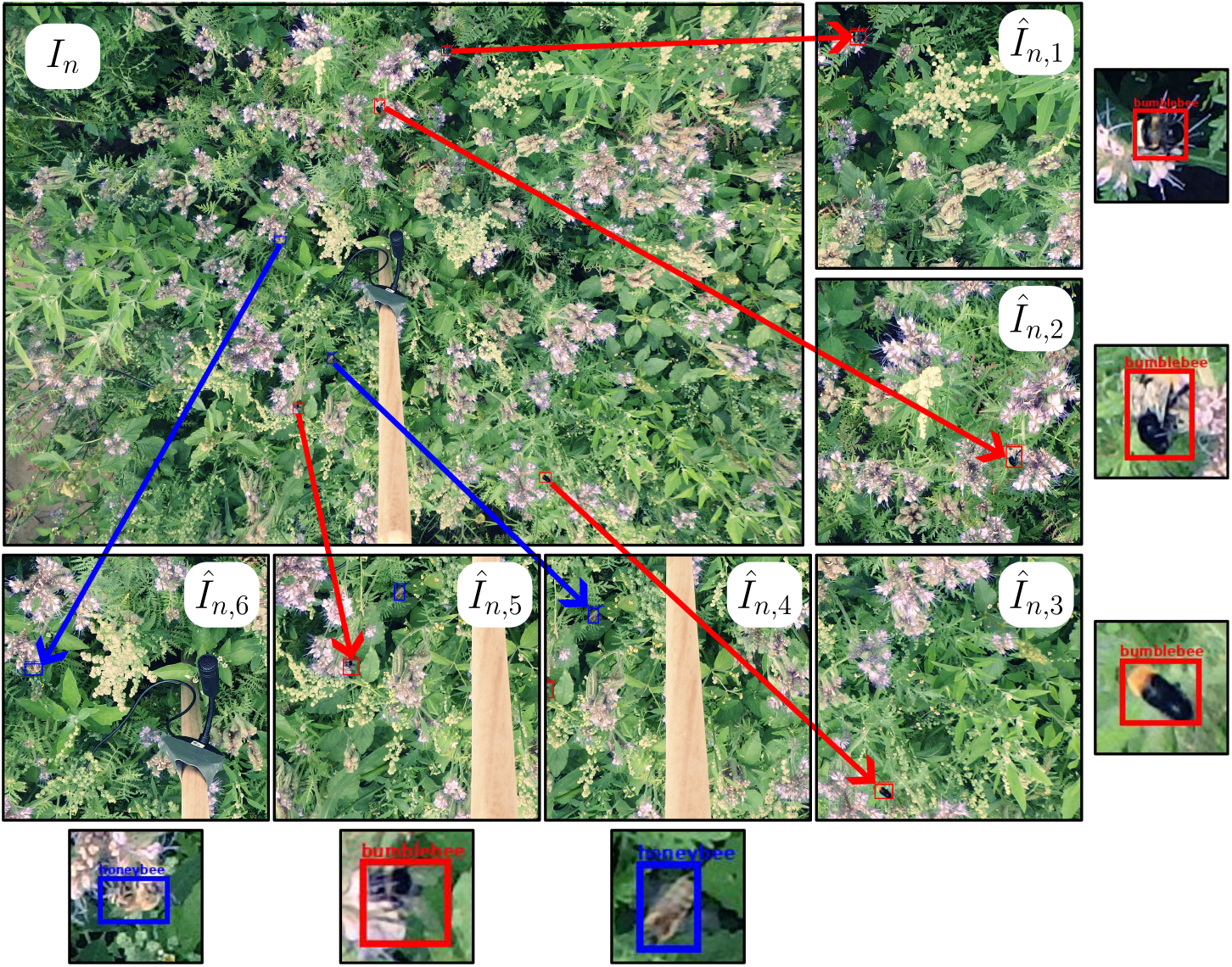
Preprocessing of images for YOLO training. An example of a camera image *I*_*n*_ cropped into smaller images 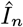 according to the prior detections for training. The honeybees are shown in blue, and the bumblebees are shown in red. We show the target pollinator in each cropped image for a clearer view.

**Fig. 4.**
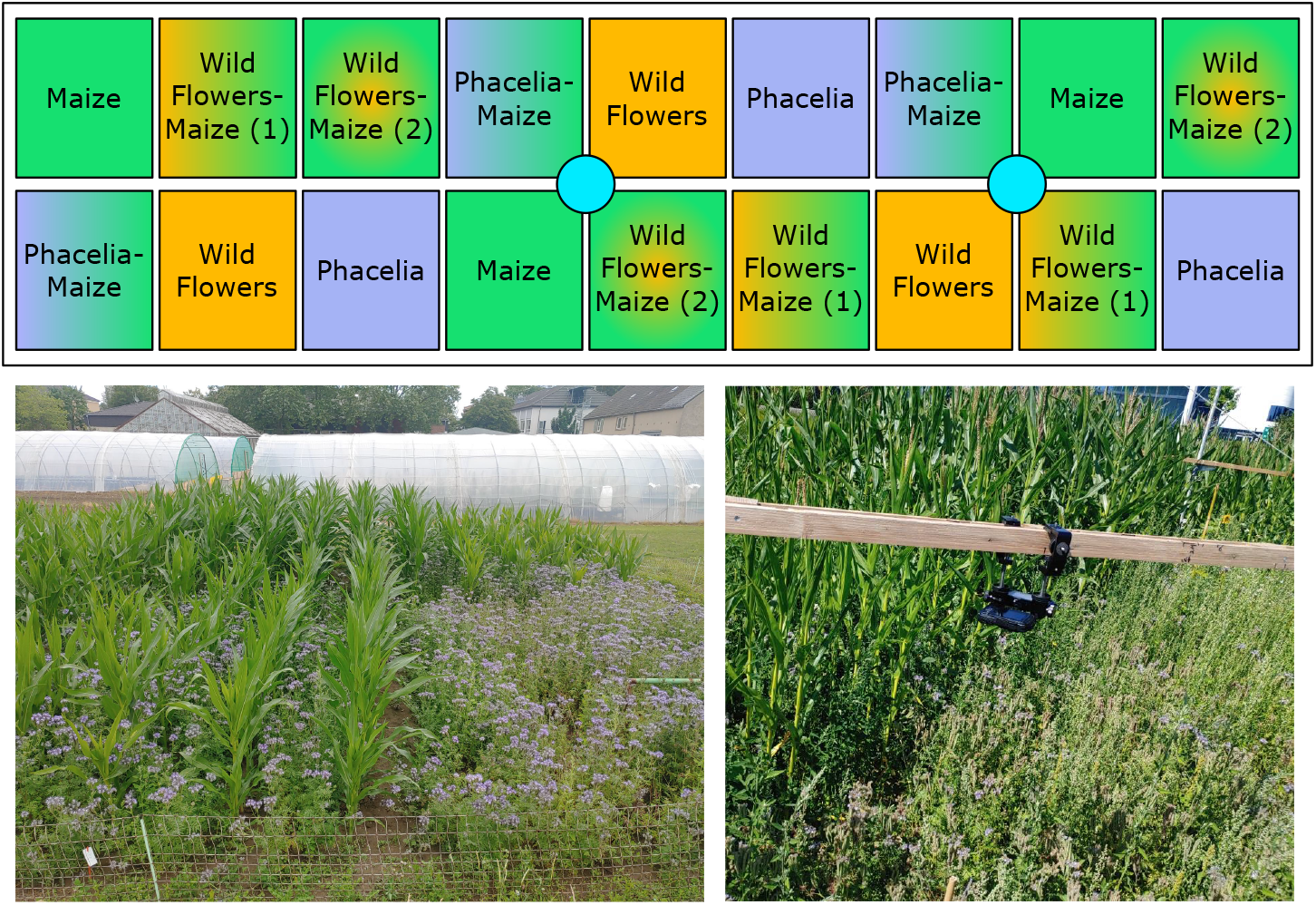
Field design of the field trial. The blue circles indicate the positions of the camera setups, each with one camera. The setups can rotate to capture data from multiple plots. Bottom left: picture of the field trial. Bottom right: picture of the camera setup.

#### Field Experiment

The experimental field (about 11 m × 38 m) in Germany with 18 plots of equal size using the randomized complete block design, as shown in Fig. 4. The area of each plot is 20 m^2^ (4 m × 5 m) with 50 cm distance between the plots. There are five distinct crop treatments in the field trial: (1) maize mono-crops, (2) phacelia flower mono-crops, (3) wildflower mixtures, (4) maize-phacelia intercrops, and (5) maize-wildflower intercrops. There are two variants of the maize-wildflowers intercrops with varying seed densities of wildflowers; refer to the supplementary material for agronomic management and design details. We capture images of all crop treatments except for the maize monocrops, where we observe only a few pollinators.

#### Camera Setups

In 2022, two camera setups, as shown in Fig. 4, automatically capture large amounts of images over different crop treatments and multiple days, with minimal manual intervention. The camera setups capture image data with consistent vantage points and minimal manual fieldwork.

Above the crop treatments, we mount a RICOH WG-70 camera on a custom camera stand. The camera triggers automatically at a predefined frequency of 0.1 Hz. We record data at each plot for at least 15 minutes. The images have a resolution of 4608 *×* 3456 pixels, resulting in a ground sampling distance of approximately 0.5 mm. The custom camera stands have adjustable heights (approximately 1.7 m above the ground) since the plant heights change throughout the season. The camera stands also had a movable cantilever arm to rotate the camera to view different crop treatments. The camera stands had a two-pronged base, which reduces the disturbance to the plots while providing a stable base. Each camera stand also features an extended arm to prevent capturing the plot edges or shadows within the camera’s field of view. These long arms are kept parallel to the ground with a supporting pulley system at the end of the cantilever arm.

When the flowers were in bloom, we collected data between 11 AM and 3 PM on five days (July 10, 17, 19, 20, and 21) in 2022. In total, we captured 2503 images for our dataset. In addition to the images from the field trial in 2022, we include images of 4032 *×*3024 pixels captured using a mobile phone with different camera angles in the phacelia mono-crops from 2021.

### 3.3 Dataset Statistics

From the annotations of the final iteration, we form a dataset for pollinator detection with accurate annotations. For the development of deep learning methods, we randomly split our dataset into train/validation/test sets. Tab. 3 shows an overview of the number of images and annotations in each split.

**Table 3.** Dataset statistics of the provided splits.

| Split | #images | #honeybees | #bumblebees | #other insects |
| --- | --- | --- | --- | --- |
| Train | 1792 | 795 | 697 | 161 |
| Validation | 384 | 120 | 133 | 36 |
| Test | 385 | 147 | 144 | 44 |
| Total | 2561 | 1062 | 974 | 241 |

Our dataset comprises data from four different flower treatments collected on multiple dates; refer to the supplementary materials for the number of images, the counts by object class, and the distribution of annotation locations in the images ℐ from 2022. We record the images for our dataset under real field conditions to capture a variety of domains across field treatments and weather conditions. In addition, we introduce 58 additional annotated images we captured using a smartphone in 2021, featuring various camera angles for future work on domain shifts.

## 4 Annotations with Varying Accuracy

### 4.1 Annotation Statistics

From each iteration *i* of the semi-automatic annotation approach, we obtain a new set of annotations ^*i*^ with decreasing errors as 𝒟 *i* increases. The train/validation/test splits for these sets of inaccurate or incomplete annotations follow that of the final dataset. Tab. 4 shows the number of pollinators we annotated after each iteration *i*, where *i* = 0 refers to the initial manual annotation. The total number of individuals increases with each iteration, except in the fifth iteration, where it decreases by 15. In *i* = 5, the number of individuals in the “other insects” class increases with respect to previous iterations; we suppose that this is because the annotations are closer to completion, and the human annotators focus more on correcting misclassification errors. The iterative semi-automatic annotation approach enhances annotation by reducing the number of insects that human annotators would have otherwise missed.

**Table 4.** Total number of annotated individuals for the dataset from each iteration.

| $i$ | #honeybees | #bumblebees | #other | #total |
| --- | --- | --- | --- | --- |
| 0 | 401 | 516 | 302 | 1219 |
| 1 | 729 | 773 | 374 | 1876 |
| 2 | 923 | 960 | 142 | 2025 |
| 3 | 1102 | 1035 | 75 | 2212 |
| 4 | 1135 | 1023 | 134 | 2292 |
| 5 (final) | 1062 | 974 | 241 | 2277 |

### 4.2 Performance Drop Caused by Erroneous Training Data

The data annotation quality should influence the model performance. To quantify the impact of annotation errors on training, we evaluate the performance of YOLO trained with each iteration’s annotations. During testing, we use the default YOLO hyperparameters, with a confidence threshold of 0.001, a non-maximum suppression IoU threshold of 0.6, and a maximum of 300 detections per image and a maximum of 200 epochs, with early-stopping after 20 epochs with no performance improvement over the validation set. Tab. 5 reports the average precision (AP_50_) and mean average precision (mAP_50_) with the IoU threshold of 50% of each YOLO weights on the test set. Unsurprisingly, the YOLO from the final iteration had the best mAP_50_, with the final iteration (*i* = 5) yielding an increase in mAP_50_ of 28.7% compared to the manual annotations (*i* = 0). While the mAP_50_ increases for intermediate iterations *i* = 0 to *i* = 2, the mAP_50_ for the third and fourth iterations are lower due to their poor performance in the “other insects” class, which may be explained by the class imbalance particularly notable for *i* = 3 and *i* = 4, as shown in Tab. 4.

**Table 5.** AP_50_ and mAP_50_ of YOLO that we train on each iteration and evaluate on the final test set. Higher values equate to better performance. The best-performing model is shown in bold.

| Iteration | AP<sub>50</sub> (%) |  |  | mAP<sub>50</sub> (%) |
| --- | --- | --- | --- | --- |
|  | honeybee | bumblebee | others |  |
| 0 | 49.0 | 70.6 | 4.4 | 41.3 |
| 1 | 74.6 | 77.8 | 10.2 | 54.2 |
| 2 | 76.1 | 86.4 | 19.5 | 60.7 |
| 3 | **83.5** | 86.7 | 5.45 | 58.5 |
| 4 | 80.6 | 87.6 | 7.73 | 58.6 |
| 5 (final) | 79.9 | **88.1** | **42.0** | **70.0** |

## 5 Baselines

To facilitate future work on pollinator monitoring, we provide several baselines for our dataset. We train and evaluate various off-the-shelf object detection methods [11,12,24] on our final dataset from *i* = 5. Refer to the supplementary material for further details on the training of each baseline. Tab. 6 reports the mAP_50_ of different object detection methods we evaluate with the final test set.

**Table 6.**
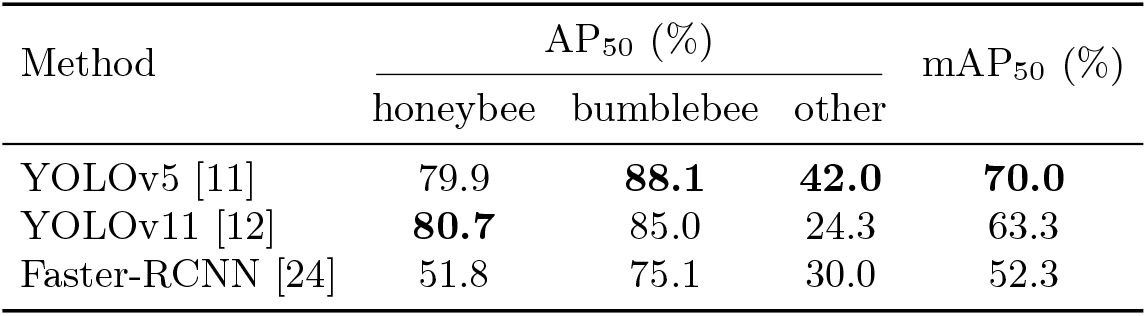
AP_50_ and mAP_50_ of object detection methods on test set annotations from different iterations. The best performance for each iteration is shown in bold.

To evaluate the methods, we report the AP_50_ and mAP_50_ for each class at an IoU threshold of 50%. Across all methods, the AP_50_ for the “other insects” class is lower than that of the honeybees and bumblebees, corresponding with the class imbalance in annotations. Additionally, this class exhibits high variance due to the presence of insects from different species, with few examples of each species. Interestingly, the YOLOv5 [11] method performs slightly better than the YOLOv11 [12], particularly with the “other insects” class. Note that we did not perform hyperparameter optimization for the baselines YOLOv11 and Faster-RCNN, and we use the hyperparameters from Section 3.1 for YOLOv5. We refer to the supplementary material for the qualitative results.

## 6 Discussion and Future Work

In this paper, we provide a dataset for pollinator detection, which we annotate via an iterative semi-automatic approach, to obtain more complete and accurate annotations by progressively correcting small object detection annotations where manual annotation is erroneous. We demonstrate that the initial manual annotations contain errors, which negatively impact model performance. However, when after we correct these annotations using an iterative semi-automatic approach, the resultant model’s performance improves. While we apply this approach specifically for pollinator detection, the iterative annotation approach can also be useful for developing other small object detection methods with sparse annotations, such as in cell detection [2] or aerial wildlife detection [25]. Moreover, with the sets of incomplete annotations that we provide, one can explore methods for improving robustness against errors in varying degrees of training data. Our dataset can also provide a means to explore data-centric AI methods toward label correction of human errors in annotations.

## 7 Conclusion

In this paper, we present a dataset for pollinator detection, focusing on honeybees and bumblebees, in various flower treatments and over multiple days. The dataset has accurate and complete annotations, which we generated through our iterative semi-automatic annotation approach that leverages YOLO to assist manual annotation and progressively refine the annotations. This iterative annotation approach also generates intermediate sets of annotations of progressively decreasing error levels and increasing number of annotations with each iteration. We quantify the performance of models trained on the intermediate annotations from each iteration, showing that the model performance improved after the corrections relative to the manual annotations. We also provide baselines for the object detection task using our dataset. Beyond enabling automatic pollinator monitoring, the iterative semi-automatic annotation approach can be useful for annotating small objects. Future work can utilize the included sets of multi-level error annotations for label correction or the development of more robust methods. This dataset supports development of deep learning methods towards automated large-scale pollinator detection under real-world field conditions.

## 8 Data Availability

Our code and dataset is made available via https://github.com/yuelinn/bee_detection.

## References

1. Amarathunga, D.C., Ratnayake, M.N., Grundy, J., Dorin, A.: Fine-grained image classification of microscopic insect pest species: Western flower thrips and plague thrips. Computers and Electronics in Agriculture 203, 107462 (2022)

2. Bertram, C.A., Aubreville, M., Marzahl, C., Maier, A., Klopfleisch, R.: A large-scale dataset for mitotic figure assessment on whole slide images of canine cutaneous mast cell tumor. Scientific Data 6(1), 274 (2019)

3. Bjerge, K., Mann, H.M.R., Høye, T.T.: Real-time insect tracking and monitoring with computer vision and deep learning. Remote Sensing in Ecology and Conservation 8(3), 315–327 (2022)

4. Blaauw, B.R., Isaacs, R.: Flower plantings increase wild bee abundance and the pollination services provided to a pollination-dependent crop. Journal of Applied Ecology 51(4), 890–898 (2014)

5. Chagnon, M., Kreutzweiser, D., Mitchell, E., Morrissey, C.A., Noome, D.A. P.J.,, der Sluijs, V.: Risks of large-scale use of systemic insecticides to ecosystem functioning and services. Environmental Science and Pollution Research 22(1), 119–134 (2015)

6. Dudley, N., Alexander, S.: Agriculture and biodiversity: a review. Biodiversity 18(2-3), 45–49 (2017)

7. Hallmann, C., Sorg, M., Jongejans, E., Siepel, H., Hofland, N., Schwan, H., Stenmans, W., Mueller, A., Sumser, H., Hoerren, T., Goulson, D., de Kroon, H.: More than 75 percent decline over 27 years in total flying insect biomass in protected areas. PLoS ONE 12(10), 1–21 (2017)

8. Herde, M., Huseljic, D., Rauch, L., Sick, B.: dopanim: A dataset of doppelganger animals with noisy annotations from multiple humans. Proc. of the Conf. on Neural Information Processing Systems (NeurIPS) 37, 51085–51117 (2024)

9. Hu, J., Yuanyuan, H., Chen, Y., Wang, H., Yasuhara, M.: Noisy ostracods: A fine-grained, imbalanced real-world dataset for benchmarking robust machine learning and label correction methods. Proc. of the Conf. on Neural Information Processing Systems (NeurIPS) 37, 50750–50771 (2024)

10. Høye, T.T., Ärje, J., Bjerge, K., Hansen, O.L.P., Iosifidis, A., Leese, F., Mann, H.M.R., Meissner, K., Melvad, C., Raitoharju, J.: Deep learning and computer vision will transform entomology. National Academy of Sciences 118(2), e2002545117 (2021)

11. Jocher, G., Chaurasia, A., Stoken, A., Borovec, J., NanoCode012, Kwon, Y., TaoXie Michael, K., Fang, J., imyhxy, Lorna, Wong, C., Yifu, Z., V, A., Montes, D., Wang, Z., Fati, C., Nadar, J., Laughing, UnglvKitDe, tkianai, yxNONG, Skalski, P., Hogan, A., Strobel, M., Jain, M., Mammana, L., xylieong: Ultralytics YOLOv5 (2022)

12. Jocher, G., Qiu, J.: Ultralytics YOLO11 (2024). https://github.com/ultralytics/ultralytics

13. Jönsson, A.M., Ekroos, J., Dänhardt, J., Andersson, G.K., Olsson, O., Smith, H.G.: Sown flower strips in southern sweden increase abundances of wild bees and hoverflies in the wider landscape. Biological Conservation 184, 51–58 (2015)

14. Kujur, V., Bedi, A.K., Saini, M.: Monitoring pollination by honeybee using computer vision. In: Proc. of Intelligent Human Computer Interaction (IHCI) (2023)

15. Li, J., Xiong, C., Hoi, S.C.: Learning From Noisy Data With Robust Representation Learning. In: Proc. of the IEEE/CVF Intl. Conf. on Computer Vision (ICCV) (2021)

16. Lin, T., Maire, M., Belongie, S., Hays, J., Perona, P., Ramanan, D., Dollár, P., Zitnick, C.L.: Microsoft COCO: Common Objects in Context. In: Proc. of the Europ. Conf. on Computer Vision (ECCV) (2014)

17. Marzahl, C., Hill, J., Stayt, J., Bienzle, D., Welker, L., Wilm, F., Voigt, J., Aubreville, M., Maier, A., Klopfleisch, R., et al.: Inter-species cell detection-datasets on pulmonary hemosiderophages in equine, human and feline specimens. Scientific Data 9(1), 269 (2022)

18. Montgomery, G.A., Belitz, M.W., Guralnick, R.P., Tingley, M.W.: Standards and best practices for monitoring and benchmarking insects. Frontiers in Ecology and Evolution 8 (2021)

19. Nguyen, D.T., Nguyen, D.M., Pham, D.T., Than, K., Pham, H.T., Vu, H.: Bayesian method for bee counting with noise-labeled data. In: Proc. of the Intl. Symp. on Information and Communication Technology (SOICT) (2023)

20. Niu, L., Tang, Q., Veeraraghavan, A., Sabharwal, A.: Learning From Noisy Web Data With Category-Level Supervision. In: Proc. of the IEEE/CVF Conf. on Computer Vision and Pattern Recognition (CVPR) (2018)

21. Pavoni, G., Corsini, M., Ponchio, F., Muntoni, A., Edwards, C., Pedersen, N., Sandin, S., Cignoni, P.: Taglab: Ai-assisted annotation for the fast and accurate semantic segmentation of coral reef orthoimages. Journal of Field Robotics (JFR) 39(3), 246–262 (2022)

22. Ratnayake, M.N., Dyer, A.G., Amarathunga, D.C., Zaman, A., Dorin, A.: Spatial monitoring and insect behavioural analysis using computer vision for precision pollination. Intl. Journal of Computer Vision (IJCV) 131(3), 591–606 (2023)

23. Ratnayake, M.N., Dyer, A.G., Dorin, A.: Tracking individual honeybees among wildflower clusters with computer vision-facilitated pollinator monitoring. PLoS ONE 16(2), 1–20 (2021)

24. Ren, S., He, K., Girshick, R., Sun, J.: Faster R-CNN: Towards real-time object detection with region proposal networks. In: Proc. of the Conf. Neural Information Processing Systems (NIPS) (2015)

25. Samiappan, S., Krishnan, B.S., Dehart, D., Jones, L.R., Elmore, J.A., Evans, K.O., Iglay, R.B.: Aerial wildlife image repository for animal monitoring with drones in the age of artificial intelligence. The Journal of Biological Databases and Curation 2024, baae070 (2024)

26. Scharr, H., Minervini, M., Fischbach, A., Tsaftaris, S.A.: Annotated Image Datasets of Rosette Plants. In: Proc. of the Europ. Conf. on Computer Vision (ECCV) (2014)

27. Sledevič, T., Matuzevičius, D.: Labeled dataset for bee detection and direction estimation on entrance to beehive. Data in Brief 52, 110060 (2024)

28. Tannous, M., Stefanini, C., Romano, D.: A Deep-Learning-Based Detection Approach for the Identification of Insect Species of Economic Importance. Insects 14(2), 148 (2023)

29. Tu, Y., Zhang, B., Li, Y., Liu, L., Li, J., Wang, Y., Wang, C., Zhao, C.R.: Learning From Noisy Labels With Decoupled Meta Label Purifier. In: Proc. of the IEEE/CVF Conf. on Computer Vision and Pattern Recognition (CVPR) (2023)

30. van der Sluijs, J.P.: Insect decline, an emerging global environmental risk. Current Opinion in Environmental Sustainability 46, 39–42 (2020)

31. Weyler, J., Magistri, F., Marks, E., Chong, Y.L., Sodano, M., Roggiolani, G., Chebrolu, N., Stachniss, C., Behley, J.: PhenoBench — A Large Dataset and Benchmarks for Semantic Image Interpretation in the Agricultural Domain. IEEE Trans. on Pattern Analysis and Machine Intelligence (T-PAMI) 46(12), 9583–9594 (2024)

32. Wu, X., Zhan, C., Lai, Y., Cheng, M., Yang, J.: IP102: A Large-Scale Benchmark Dataset for Insect Pest Recognition. In: Proc. of the IEEE/CVF Conf. on Computer Vision and Pattern Recognition (CVPR) (2019)

33. Ziesche, T.M., Ordon, F., Schliephake, E., Will, T.: Long-term data in agricultural landscapes indicate that insect decline promotes pests well adapted to environmental changes. Journal of Pest Science (2023)

