## Supplement for "A Computer Vision Dataset for Pollinator Detection under Real Field Conditions"

### Supplementary Material

In this supplementary material, we provide additional details about our proposed approach.

#### A Field Design and Agronomic Management

Table 1. Experimental key treatments analyzed in this study in 2022.

| Treatment ID | Species/seeds |
|---|---|
| Maize | Maize (75 cm row distance, 9 plants per $m^2$) |
| Phacelia | Phacelia |
| Wildflowers | Mixed flowers (phacelia and other species) |
| Phacelia-Maize | Phacelia and maize (75 cm maize row distance, 9 plants per $m^2$) |
| Wildflowers-Maize | Mixed flowers (phacelia and other species) and maize (75 cm maize row distance) |

The field design in 2022 comprised of both monoculture maize and flowering plants-only treatments (with and without maize). Table 1 shows the details of the five treatments, each with three replicates, but the pollinators were monitored only in selected treatments with flowers.

Maize (*Zea mays*, varieties Rudint in 2021 and Rubesto ÖKO in 2022) was sown manually about 5 cm deep with a row spacing of 75 cm (9 plants per $m^2$) with five rows per plot. Phacelia (*Phacelia tanacetifoloa*, variety Boratus) monocrops was sown solely with 1 g per $m^2$. In the phacelia-maize and wildflower-maize intercrop treatments, between each row of maize, two rows of flowering plants were sown. Thus, flowering plants covered about 25 % of the area in the intercrops, while the maize was grown at 100 % of sowing density equal to that of its corresponding monoculture treatment, resulting in additive design [5], which is also sometimes called an intermediate mixture design, where the summed relative crop densities are above 100% but below 200 % [1].

In 2022, we sowed the commercially available flower strip seeds OPTIMA® WildLife BIENENWEIDE SCHLESWIG-HOLSTEIN (referred to as "wildlife bee") for treatments F and FM consisting of the species phacelia, buckwheat (*F. esculentum Moench*), flax (*Linum usitatissimum*), sunflower (*Helianthus annuus*), mallow (*Malva sylvestris*), dill (*Anethum graveolens*), crimson clover (*Trifolium incarnatum*), false flax (*Camelina sativa*), serradella (*Ornithopus sativus*), spring vetch (*Vicia lathyroides*), Persian clover (*Trifolium resupinatum*), Alexandrine clover (*Trifolium alexandrinum*), and marigold (*Tagetes erecta*) with a sowing density of 1 g per $m^2$. Sowing and the final harvest took place on May 5, May 6, and September 8, 2022. Manual mechanical weeding was conducted on June 1 and June 14, 2022. NPK fertilizers were applied on June 9 (50 kg $ha^{-1}$ of N) in treatments P, F and M and 100 kg $ha^{-1}$ of N in treatments PM and FM. Table 2 shows the weather data during the data collection.

Table 2. Mean monthly air temperature and monthly precipitation sums for April to September 2021 and 2022 at the field site.

| Months | Temperature | Precipitation |
|---|---|---|
| April | 10.4 | 45.1 |
| May | 16.6 | 55.3 |
| June | 19.6 | 89.8 |
| July | 21.1 | 18.9 |
| August | 22.6 | 8.6 |
| September | 20.7 | 19.9 |

Table 3. Number of pollinators detected (absolute value) on given days and treatments in 2022. The intercropping of phacelia-maize treatment is denoted with the code 'PM', mixed flowers treatment is denoted with the code 'F', sole crop phacelia treatment is denoted with the code 'P', and intercropping of mixed flowers-maize treatment is denoted with the code 'FM'.

| Date | #honeybees | | | | #bumblebees | | | | #other | | | | Total #individuals | | | |
|---|---|---|---|---|---|---|---|---|---|---|---|---|---|---|---|---|
| | PM | F | P | FM | PM | F | P | FM | PM | F | P | FM | PM | F | P | FM |
| July 10 | 119 | 20 | 93 | 18 | 104 | 103 | 256 | 9 | 3 | 14 | 36 | 6 | 226 | 137 | 385 | 33 |
| July 17 | 34 | 12 | 60 | 20 | 82 | 19 | 45 | 11 | 11 | 12 | 9 | 6 | 127 | 43 | 114 | 37 |
| July 19 | 52 | 50 | 33 | 3 | 23 | 3 | 8 | 5 | 9 | 14 | 9 | 1 | 84 | 67 | 50 | 9 |
| July 20 | 41 | 42 | 27 | 6 | 24 | 33 | 23 | 28 | 12 | 8 | 12 | 5 | 77 | 83 | 62 | 39 |
| July 21 | 1 | 2 | 57 | 15 | 59 | 26 | 10 | 2 | 0 | 2 | 7 | 24 | 60 | 30 | 74 | 41 |
| Total | 247 | 126 | 270 | 62 | 292 | 184 | 342 | 55 | 35 | 50 | 73 | 42 | 574 | 360 | 685 | 159 |

#### B Dataset Statistics

Table 3 shows the distribution of annotations for each treatment and date and Table 4 shows the number of images captured per date Fig. 1 shows the distribution of the objects in $\mathcal{I}$. The distribution is denser at the center since the images were centered around the flowers in the field; but the corners of the image also contain a fair amount of annotated pollinators.

#### C Baseline Training Details

##### C.1 YOLOv11

We trained a YOLOv11 [3], specifically the YOLOv11m architecture, initialized with weights pre-trained on COCO, for the maximum of 200 epochs and early-stopping after 20 epochs when the performance metrics on the validation set does not improve. We used the constant learning rate of 0.0014 with the AdamW optimizer with momentum of 0.9, batch size of 4, image size of 3600, and also

Table 4. Number of images captured on given days and treatments in 2022.

| Date | phacelia-maize | wildflowers | phacelia | wildflowers-maize |
|---|---|---|---|---|
| July 10 | 104 | 116 | 109 | 105 |
| July 17 | 107 | 112 | 120 | 105 |
| July 19 | 108 | 226 | 109 | 104 |
| July 20 | 120 | 106 | 107 | 111 |
| July 21 | 109 | 160 | 169 | 196 |

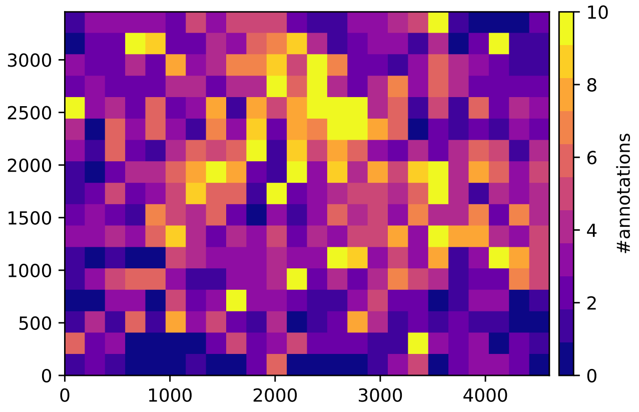


Fig. 1. Distribution of annotation locations. Dark purple indicates lower annotation count, while bright yellow indicates higher annotation counts. The distribution of annotations covers most of the image, with more annotations towards the center.

applied augmentations of color jitter in HSV space, translation, resizing, flipping, and mosaic.

##### C.2 Faster-RCNN

We trained a Faster-RCCN[4] with a ResNet-50 a backbone, pre-trained on the COCO dataset, for 50 epochs with a batch size of 2, using Adam for optimization, and at the maximum image size of 4608. We set the initial learning rate to 0.0001 and apply an exponentially decaying schedule. We use the weights with the best mAP performance at IoU threshold steps from 50% to 95% on the validation set.

#### D Qualitative Pollinator Detection Results

Fig. 2 shows the qualitative results of each baseline methods, with predictions with confidence of at least 0.5. The images were cropped to better show the detected pollinators.

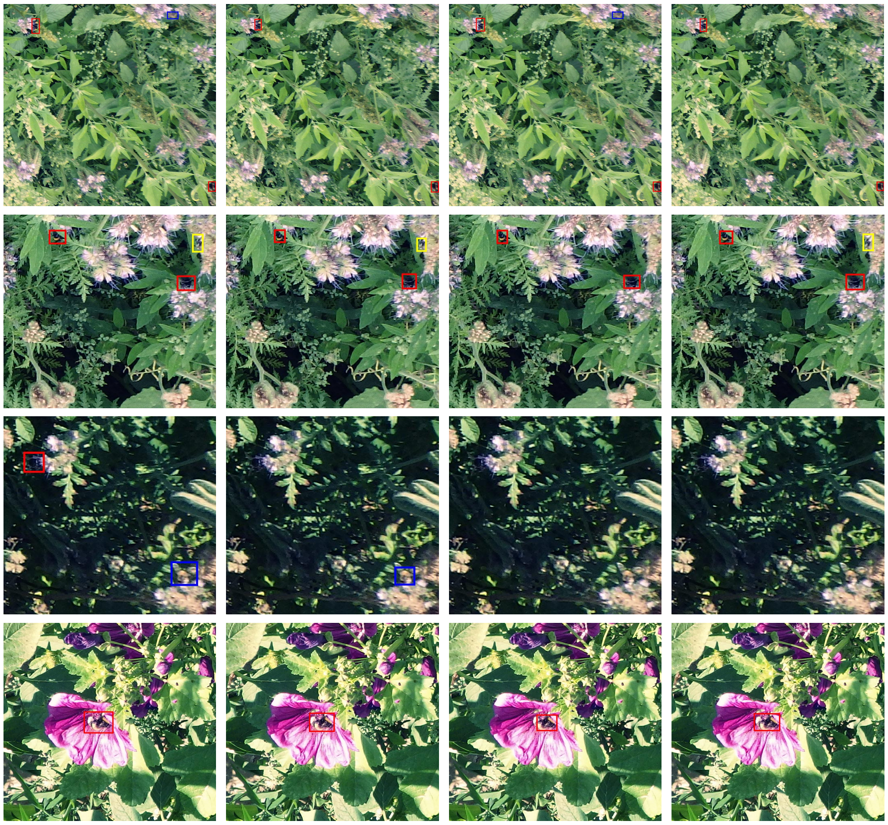

(a) Ground Truth (b) YOLOv5 (c) YOLOv11 (d) Faster-RCNN

Fig. 2. Sample qualitative results from the test set with predictions from YOLOv5 [2], YOLOv11 [3], and Faster-RCNN[4]. The honeybees are marked in blue, bumblebees in red, and other bees in yellow.

#### References


1. Demie, D.T., Döring, T.F., Finckh, M.R., van der Werf, W., Enjalbert, J., Seidel, S.J.: Mixture × genotype effects in cereal/legume intercropping. Frontiers in Plant Science **13** (2022) 1
2. Jocher, G., Chaurasia, A., Stoken, A., Borovec, J., NanoCode012, Kwon, Y., TaoXie, Michael, K., Fang, J., imyhxy, Lorna, Wong, C., Yifu, Z., V, A., Montes, D., Wang, Z., Fati, C., Nadar, J., Laughing, UnglvKitDe, tkianai, yxNONG, Skalski, P., Hogan, A., Strobel, M., Jain, M., Mammana, L., xylieong: Ultralytics YOLOv5 (2022) 4
3. Jocher, G., Qiu, J.: Ultralytics YOLO11 (2024). `https://github.com/ultralytics/ultralytics` 2, 4
4. Ren, S., He, K., Girshick, R., Sun, J.: Faster R-CNN: Towards real-time object detection with region proposal networks. In: Proc. of the Conf. Neural Information Processing Systems (NIPS) (2015) 3, 4
5. Zustovi, R., Landschoot, S., Dewitte, K., Verlinden, G., Dubey, R., Maenhout, S., Haesaert, G.: Intercropping indices evaluation on grain legume-small grain cereals

mixture: A critical meta-analysis review. Agronomy for Sustainable Development **44**(1), 5 (2024) 1
